# Structural basis for the concerted antiphage activity in the SIR2-HerA system

**DOI:** 10.1101/2023.11.13.566805

**Authors:** Guimei Yu, Fumeng Liao, Chendi Zhang, Xuzichao Li, Qiuqiu He, Hang Yin, Zhuang Li, Heng Zhang

## Abstract

Recently, a novel two-gene bacterial defense system against phages, encoding a SIR2 NADase and a HerA translocase, has been identified. However, the molecular mechanism of the bacterial SIR2-HerA immune system remains unclear. Here, we determine the cryo-EM structures of SIR2, HerA and their complex in different functional states. The SIR2 proteins oligomerize into a dodecameric ring-shaped structure consisting of two layers of interlocked hexamers, in which each SIR2 unit exhibits an auto-inhibited conformation. Distinct from the canonical AAA+ proteins, the HerA hexamer in this antiphage system adopts a split spiral arrangement, resembling the substrate-binding state, which is stabilized by a unique C-terminal extension. SIR2 and HerA proteins assemble into a ∼ 1.1 MDa torch-shaped complex to fight against phage infection. Importantly, disruption of the interactions between SIR2 and HerA largely abolishes the antiphage activity. Interestingly, HerA binding alters the oligomer state of SIR2, switching from a 12-mer state to a 14-mer state. On the other hand, binding of SIR2 stimulates the ATPase activity of HerA. Together, our study not only provides a structural basis for the functional communications between SIR2 and HerA proteins, but also unravels a novel concerted antiviral mechanism through nucleotide (NAD^+^ and ATP) depletion.

## Introduction

Bacteria and archaea have evolved diverse antiviral systems to confront the invading mobile genetic elements (MGEs), such as bacteriophages and plasmids (Dy *et al*, 2014; Labrie *et al*, 2010; Stern & Sorek, 2011). Notably, the innate immune systems of prokaryotes combat viral infection through diverse mechanisms, for example, the restriction-modification (R-M) systems and abortive infection (Abi) systems. However, the distribution of antiviral systems is not uniform, with different bacteria or archaea encoding distinct sets of systems to counteract infections. The prokaryotic genes encoding antiviral systems are frequently located together in genomes, forming “defense islands” (Makarova *et al*, 2013; Makarova *et al*, 2011). This phenomenon has catalyzed the rapid expansion of new antiviral systems from microbial pangenomes in recent years (Doron *et al*, 2018; Gao *et al*, 2020; Millman *et al*, 2022). Dozens of new antiphage defense systems have been recently identified, wherein diverse enzymatic activities are recruited for defense responses. Despite extensive efforts, the molecular mechanisms of most new antiphage defense systems remain unclear.

SIR2 domain-containing proteins are widespread across all domains of life. While eukaryotic and archaeal SIR2 domains, together with the NAD^+^ cofactor, exert ADP ribosyl transferase and protein deacetylase activities (North & Verdin, 2004), multiple bacterial SIR2 domain-containing proteins have recently been found in various antiphage defense systems, such as prokaryotic Argonautes (pAgo), Thoeris, AVAST and defense-associated sirtuin (DSR). These proteins deplete NAD^+^ (NADase activity) to restrict phage infection (Doron *et al*., 2018; Gao *et al*., 2020; Garb *et al*, 2022; Koopal *et al*, 2022; Makarova *et al*, 2009; Zaremba *et al*, 2022). A new two-gene antiphage defense system encoding SIR2 and a HerA-like DNA helicase has recently been identified (Gao *et al*., 2020; Garb *et al*., 2022). Transformation of bacterial strains lacking the system with vectors containing SIR2-HerA led to resistance to multiple phages and NAD^+^ depletion upon infection. Catalytically dead mutation in either protein encoded by the SIR2-HerA system abrogated phage restriction, suggesting both the NADase and the ATPase/Helicase activities are required for antiphage immunity. However, the molecular mechanism underlying the HerA-dependent NADase activation is not clear.

Here, we present multiple cryo-EM structures of the SIR2-HerA system in different states (13 structures). Combined with biochemical and mutagenesis studies, we have further elucidated mechanisms underlying NADase and ATPase activation in this system. More importantly, our findings reveal that both NAD^+^ and ATP depletion, crucial for phage resistance, are modulated by the complex assembly. In summary, our studies demonstrate that SIR2 and HerA form a stable, torch-like supramolecular complex of about 1.1 MDa, potentially facilitating the efficient coupling of the NADase activity of SIR2 and the ATPase/Helicase activity of HerA in bacterial immunity.

## Results

### SIR2 assembles into a dodecamer

To elucidate the functional role of SIR2 protein in the SIR2-HerA antiphage defense system, we first expressed and purified SIR2 protein from *Paenibacillus sp*. 453MF (PsSIR2) for cryo-EM structural characterization (Figure S1A). A 3.2 Å cryo-EM map was generated, which enabled us to unambiguously build the atomic model of the PsSIR2 protein (Figure S2, Table S1). Remarkably, PsSIR2 protein oligomerizes into a ring-shaped dodecamer with a diameter of approximately 170 Å, a height of 80 Å and an inner diameter of about 50 Å (Figure 1A). Consistently, the analytical centrifugation analyses also supported the dodecameric state of PsSIR2 in solution (Figure S1B). In addition, the cryo-EM structure of SIR2 protein from the SIR2-HerA system of *Planctomycetes* HKU-PLA5 (PhpSIR2) was also resolved in the same dodecameric state (Figure S3, Table S2), suggesting a conserved oligomerization mode of SIR2 protein among the SIR2-HerA antiphage defense systems. Due to the high resolution, we mainly focused on the PsSIR2 protein for structural analysis.

**Figure 1.**
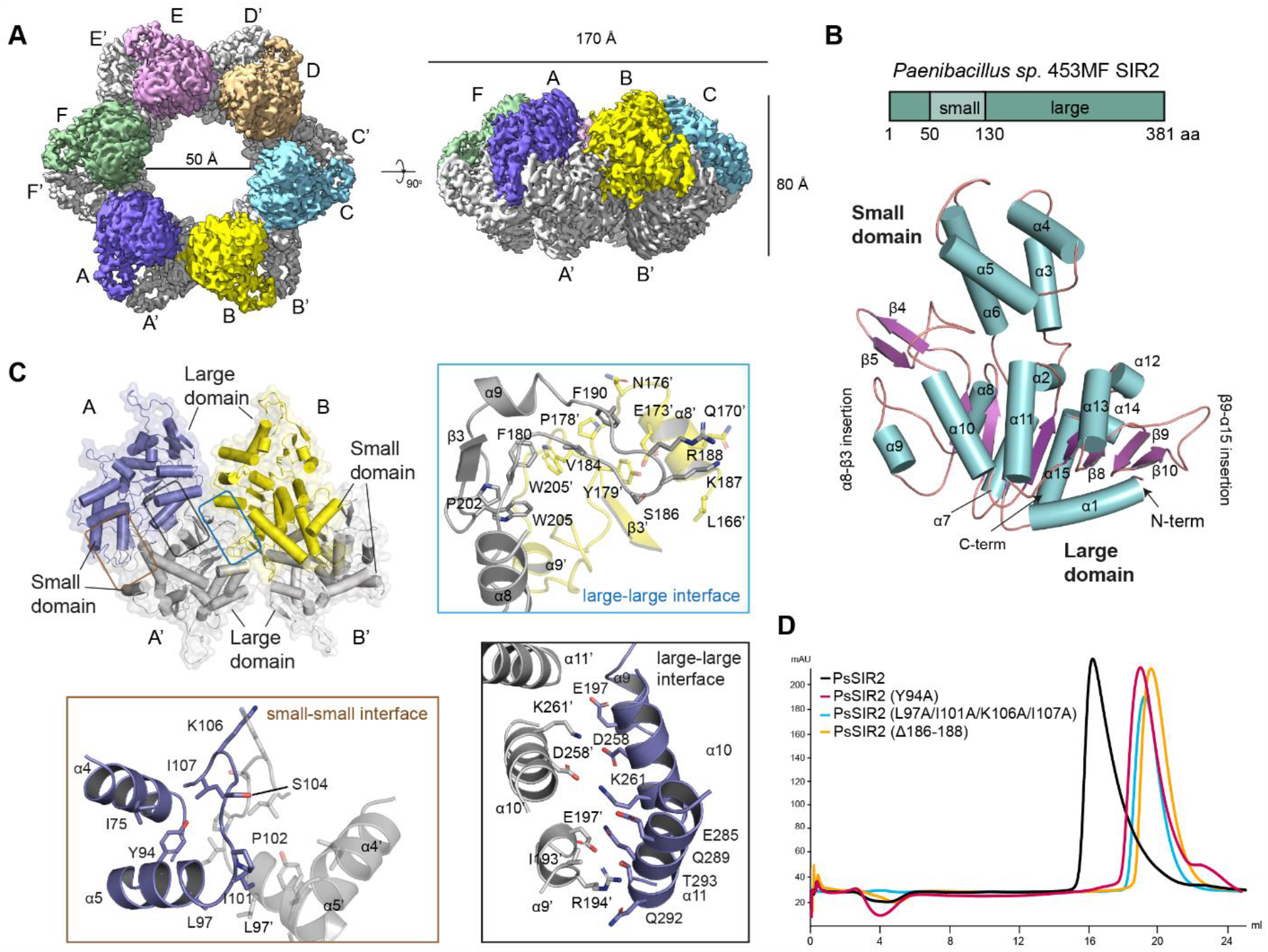
SIR2 assembles intro a ring-shaped dodecamer. (**A**) Cryo-EM map of PsSIR2 at the side (upper) and top (lower) views. The dimensions of the inner and outer diameters and the height are indicated. (**B**) Domain organization and atomic structure of PsSIR2 subunit. The large domain of PsSIR2 is composed of a central β-sheet flanked by multiple helices and is capped by a small protruding domain. The insertions were identified in comparison to archaeal and eukaryotic SIR2 proteins. (**C**) In the ring-shaped PsSIR2 dodecamer, two PsSIR2 molecules from each layer dimerize head-to-head, interact with each other through both small and large domains, establishing two major interfaces: the small-small interface (brown box) and the large-large interface (black box). The assembly of PsSIR2 dodecamer is mediated by the interactions between two large domains of neighboring dimers. The inserted panels provide a close-up view of the three interfaces, key residues at these interfaces are represented as sticks. (**D**) Gel filtration analysis of wild-type (WT) and mutant PsSIR2 proteins.

The single PsSIR2 molecule displays a bilobed structure consisting of a large domain and a small domain (Figure 1B), akin to other SIR2-containing proteins (North & Verdin, 2004). The large domain adopts a Rossmann-fold with a central eight-stranded β-sheet sandwiched by eleven helices and a β-hairpin (Figure 1B). The small domain is a protrusion from the large domain (between α2-α7), with four helices (α3-α6) forming a helical bundle. Compared with archaeal and eukaryotic SIR2 proteins like *Archaeoglobus fulgidus* SIR2-Af1 (Min *et al*, 2001) and yeast Hst2p (Zhao *et al*, 2003), two insertions (the α8-β3 insertion and the β9-α15 insertion, respectively) at the two sides of the large domain were identified (Figure 1B and S4).

The dodecameric PsSIR2 proteins consist of two layers of interlocked hexamers, in which two PsSIR2 molecules from each layer dimerize head-to-head through both the large and small domains, thereby conjoining the two layers (Figure 1C). The dimer interface of the small domains is constituted by the symmetric α5-α6 loops extending towards α4 and α5 helices *in trans* (Figure 1C). The α9-α11 helices from the large domains further stabilize the dimerization. Alanine substitution of residues in the dimer interface, such as Y94A and L97A/I101A/K106A/I107A, substantially impaired SIR2 assembly (Figure 1D). The six dimer units pack together to form the ring, with the large domain mediating the inter-dimer interaction. Particularly, the α8-β3 insertion loops swap between the large domains of two adjacent dimers, reinforcing SIR2 assembly through polar and hydrophobic interactions (Figure 1C, top). As expected, truncation of this loop (Δ186-188) altered the oligomerization state of SIR2 (Figure 1D).

### The NADase activity of SIR2 is autoinhibited in the apo state

The potential NADase catalytic pocket in the large domain is located at the interface between the large and small domains (Figure 2A). Superimposition of the catalytic pockets of PsSIR2 and SIR2-Af1 in complex with NAD^+^ revealed that the bound NAD^+^ would clash with the α2-α3 loop connecting the two domains, and α3 and α5 helices in the small domain (Figure 2B). These observations suggest that the SIR2 protein alone in the SIR2-HerA antiphage system is likely to be autoinhibited by the small domain, and conformational movement of the small domain would be required for binding to NAD^+^. Indeed, *in vitro* NADase assay showed no NAD^+^ hydrolysis by PsSIR2 protein alone (Figure 2D). However, truncation of the small domain (Δ53-130 aa) substantially restored PsSIR2 NADase activity (Figure 2D). Additionally, alanine substitutions of residues in the α15 helix (F361A/Q362A), with the potential to alter the conformation of the α2-α3 loop, also partially activated the NADase activity of PsSIR2 (Figure 2C-D). Next, we investigated whether the dodecameric assembly causes autoinhibition. The mutations that disrupt PsSIR2 dodecamer formation were tested, but only slight activation was observed (Figure 1D and S5A), suggesting that the SIR2 NADase activity was mainly inhibited within the subunit by the small domain rather than the formation of high-order assembly.

**Figure 2.**
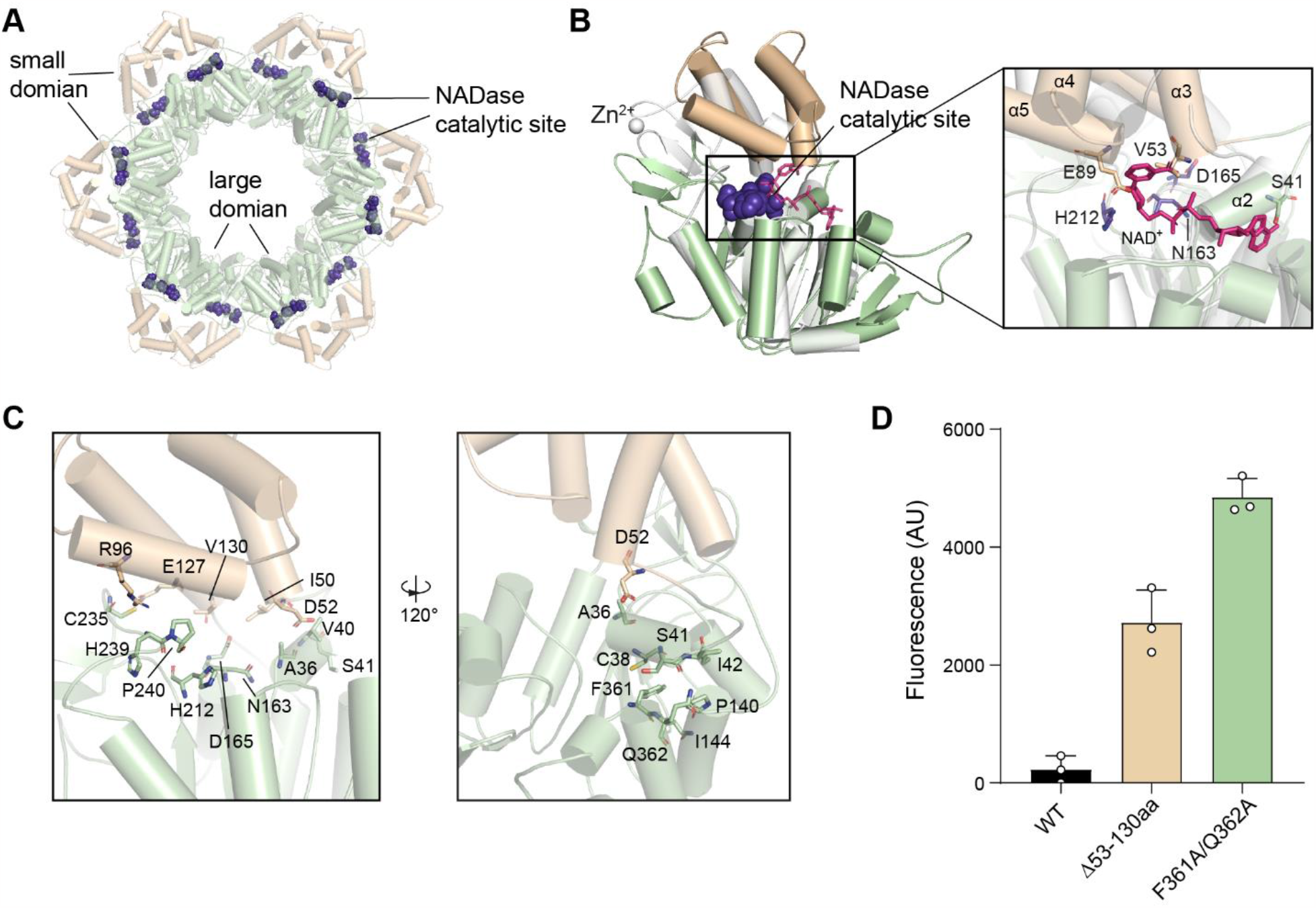
SIR2 is autoinhibited by the small domain. (**A**) The NADase catalytic pockets in PsSIR2 dodecamer. PsSIR2 is colored by domains, putative catalytic residues are shown as purple spheres. (**B**) Structural superposition of PsSIR2 and *Archaeoglobus fulgidus* sirtuins (SIR2-Af1) in complex with NAD^+^. PsSIR2 is colored by domains and the SIR2-Af1 is colored in white. The putative catalytic residues N163, D165, and H212 in PsSIR2 are shown as purple spheres. (**C**) The interaction interfaces between the small and large domains in PsSIR2 (left panel). Key residues at the interface are shown as sticks. The extensive interactions between the C-terminal distal α15 helix and α2, α7 helices in the large domain further stabilize the small-large interaction interface in the PsSIR2 (right panel). Key residues at the interfaces are represented in stick representation. (**D**) In vitro NAD^+^ degradation assay of WT and mutants of PsSIR2. The experiment was repeated three times and the data are presented as mean ± SEM.

### Cryo-EM Structures of HerA

The HerA protein in the SIR2-HerA system is a member of the AAA+ superfamily of proteins (ATPase associated with diverse cellular activities), which typically polymerizes into hexameric structures. The ATPase domain of AAA+ proteins is thought to undergo conformational changes during ATP hydrolysis, thereby generating mechanical forces necessary for substrate remodeling (Neuwald *et al*, 1999; Ogura & Wilkinson, 2001; White & Lauring, 2007). Next, we expressed and purified the HerA protein from *Paenibacillus sp*. 453MF SIR2-HerA antiphage system (PsHerA). As indicated by the analytical centrifugation analyses (Figure S5B), PsHerA behaves as a hexamer in solution, reminiscent of other AAA+ proteins. Cryo-EM structures of PsHerA in complex with adenosine 5′-O-(3-thio) triphosphate (ATPγS), a non-hydrolysable ATP analog, were determined (Figure S6, Table S1), confirming the hexamer assembly. Three subpopulations of HerA were identified, containing three, four and five ATPγS molecules, respectively (Figure S7A). The structural analysis of PsHerA primarily focuses on the state bound with five ATPγS molecules unless otherwise stated) (Figure 3A, C, D).

**Figure 3.**
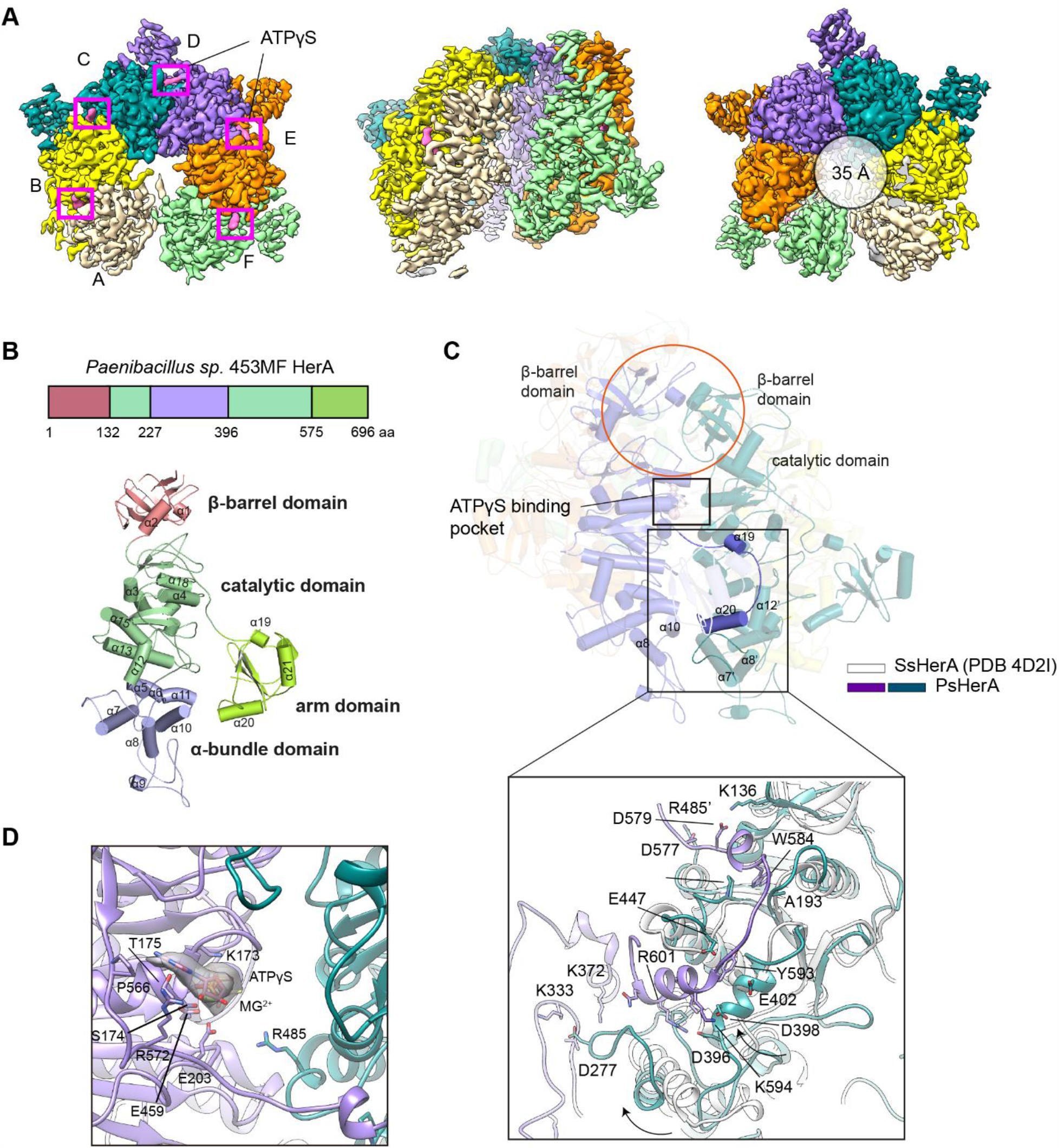
Cryo-EM structures of PsHerA. (**A**) Cryo-EM structure of PsHerA in complex with ATPγS. The top, side, and bottom views are shown, respectively. The cryo-EM densities of the ATPγS molecules are colored in pink and marked with pink rectangles. The diameter of the central channel is about 35 Å. (**B**) Domain organization and subunit structure of PsHerA. PsHerA contains four major domains, including the N-terminal β-barrel domain (salmon), the middle catalytic RecA-like domain (pale green), the bottom α-bundle domain (light blue), and the arm domain (lime) at the side. (**C**) Subunit-subunit interactions in PsHerA hexamer. The β-barrel domain associates with the groove formed by β-barrel and RecA-like domains in the neighbor subunit (salmon circle), and the arm domain further wraps the neighbor RecA-like domain (black rectangle), which together stabilize the hexamer assembly. The lower panel provides detailed insights into the interactions between the arm and RecA-like domains, key residues involved in the interactions are shown as sticks (lower panel). The cartoon model in gray represents the structure of SeHerA (PDB ID: 4D2I), which is the closest homolog of PsHerA in Dali search. (**D**) ATPγS-binding pocket in PsHerA. Cryo-EM density for bound ATPγS molecule is colored in grey. ATPγS molecule and residues in the α-bundle domain responsible for ATPγS binding are shown as sticks.

The single PsHerA molecule is composed of four domains: the N-terminal β-barrel domain (aa 1-132), the middle RecA-like AAA+ catalytic domain (aa 133-227, 397-575), the α-helix bundle domain at the bottom (aa 228-396), and the flanked C-terminal arm domain (aa 576-696) (Figure 3B). The six PsHerA subunits are arranged in a tilted configuration, forming a hexameric ring (Figure 3A). In particular, the β-barrel domain inserts into a shallow groove between the β-barrel and RecA-like catalytic domains *in trans* (Figure 3C). Additionally, the arm domain, especially the α19-20 helices and their linker, wraps around the α13, α15 and α12 helices in the catalytic domain *in trans* (Figure 3C). The nucleotide-free subunit A shows increased flexibility compared to those in the nucleotide-bound state (subunits B-F) (Figure S7B). While subunits B-E were nearly identical, a slight rotation of subunit F, particularly at the catalytic pocket, was observed, potentially representing a primed state for ATP hydrolysis in the sequential ATP hydrolysis model (Ogura & Wilkinson, 2001). Similar structural features were also evident in the structure of PsHerA in complex with AMP-PNP (Figure S7C).

### HerA adopts a split spiral conformation

A Dali search identified the HerA protein from *Sulfolobus solfataricus* (SsHerA) as the closest structural homolog to PsHerA (Rzechorzek *et al*, 2014) (Figure S7D). The substrate-free SsHerA hexamer exhibits a symmetric, closed conformation (Figure S7D). By contrast, despite the absence of a DNA substrate, the PsHerA hexamer exhibits a split spiral conformation (Figure 3A), which is distinct from the canonical substrate-free AAA+ hexamer but reminiscent of the substrate-bound state (Fernandez & Berger, 2021; Li *et al*, 2022; Puchades *et al*, 2020). Nevertheless, the partially open channel of PsHerA, with a diameter of about 35 Å, appears to be large enough to accommodate a potential DNA substrate (Figure 3A). Structural comparison with SsHerA further revealed a unique C-terminal arm domain for PsHerA (Figure 3B-C). Notably, the arm domain makes direct interactions with the α12 helix from the adjacent subunit, initiating a clockwise movement of the α-helix bundle domain (Figure 3C). This movement would enable the α-helix bundle domain to establish contacts with the neighboring subunit, thus stabilizing the spiral conformation of PsHerA. We also determined the structure of HerA from *Planctomycetes* HKU-PLA5 (PhpHerA) (Figure S9, Table S2). PhpHerA bears the same insertions and was also resolved in the same spiral conformation (Figure S8F-G), further highlighting the conserved conformation of HerA within the SIR2-HerA immune systems.

### Structures of the SIR2-HerA complex

Both SIR2 and HerA are required for phage resistance (Gao *et al*., 2020; Garb *et al*., 2022), indicating the functional interplay between SIR2 and HerA. As expected, co-elution of SIR2 and HerA proteins was observed in the gel filtration analysis (Figure S10), suggesting a direct interaction between them. Cryo-EM structures of the PsSIR2-HerA complex were then determined to further elucidate the association (Figure S11, Table S3). PsSIR2 and PsHerA proteins assemble into a ∼ 1.1 MDa torch-like complex, with the ring-shaped PsSIR2 docked over the potential DNA binding channel of PsHerA (Figure 4A).

**Figure 4.**
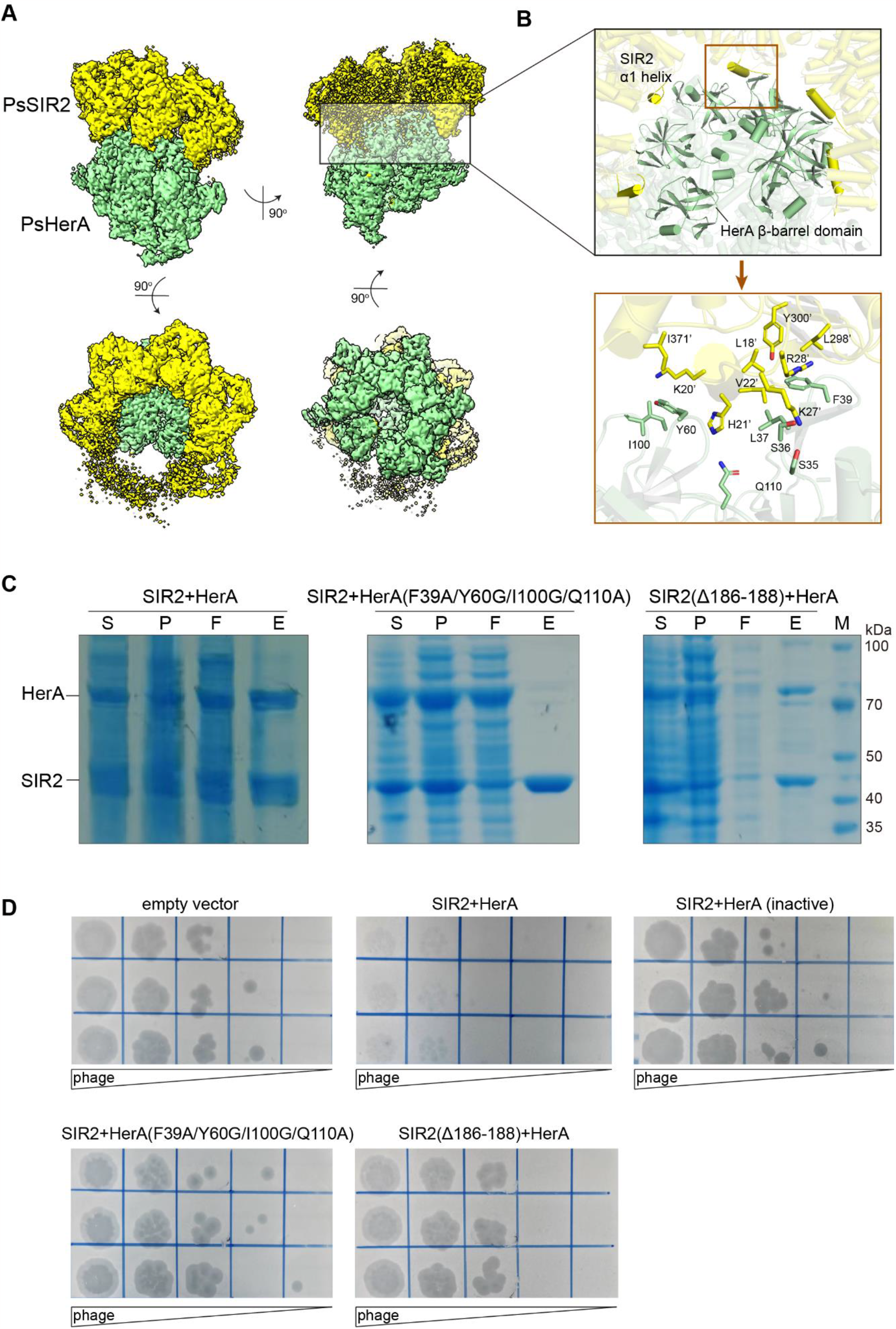
Characterization of the PsSIR2-HerA complex. (**A**) Cryo-EM structure of the PsSIR2-HerA complex. PsSIR2 proteins (yellow) in the complex are presented as 14-mer, landing at the top of PsHerA hexamer (green). (**B**) Close up views of the SIR2-HerA interacting interface. Every two β-barrel domains of HerA lock one SIR2 α1 helix (upper panel). The lower panel illustrates detailed insights into the SIR2-HerA interaction interface, with key residues at the interaction interface depicted as sticks. (**C**) Co-purification analysis of PsSIR2 and PsHerA. The 6XHis tag was sub-cloned to PsSIR2 protein in individual experiments. The co-purification assays were examined using Ni-NTA resin. S: supernatant of the cell lysis; P: pellet of cell debris; F: the flow through; E: the eluted fraction using imidazole. (**D**) Phage challenge assays with WT or mutants of PsSIR2-HerA complex. Transformation of plasmids expressing PsSIR2-HerA defense system into *E*.*coli* MG1655 substantially protected the cells from T7 phage. The protection was lost for the catalytic dead mutant. The mutants blocking the PsSIR2-HerA association or impairing PsSIR2 oligomerization failed to defend against phage infection.

PsSIR2 interacts with PsHerA primarily through the N-terminus α1 helix, which is bracketed by two β-barrel domains of the PsHerA hexamer (Figure 4B). As anticipated, substitutions of the interface residues in HerA, such as F39A/Y60A/I100G/Q110A, disrupted the PsSIR2-HerA assembly in the His-tagged pull-down assay (Figure 4B-C). To evaluate the functional importance of the SIR2-HerA interactions, we performed a bacteriophage infection plaque assay. Consistent with our *in vitro* binding results, this mutation failed to provide defense against phage (Figure 4D), supporting the notion that the SIR2-HerA assembly is essential for bacterial immunity. Notably, we found that mutation disrupting the dodecamer assembly of PsSIR2 (Δ186-188) was capable of binding PsHerA but rendered bacteria susceptible to phage infection (Figure 4C-D), indicating that the SIR2 self-association is important for the NADase activity.

### HerA specifically binds to tetradecameric SIR2

Given that two adjacent β-barrel domains of the PsHerA hexamer are responsible for recognizing one α1 helix of PsSIR2, the PsHerA hexamer would be capable of binding up to five PsSIR2 molecules. As anticipated, a portion of the ring-shaped PsSIR2 could not be clearly seen in the EM density possibly due to the absence of direct interaction with PsHerA (Figure 4A). As aforementioned, PsSIR2 alone is present as a dodecamer. However, the PsHerA hexamer is encircled by the tetradecameric rather than the dodecameric PsSIR2 in all the resolved complexes (Figure S11). To elucidate the mechanism behind the binding specificity of PsHerA, we superimposed the dodecameric PsSIR2 with the tetradecameric PsSIR2 in the PsSIR2-HerA complex (Figure 5A). While all five α1 helices of tetradecameric PsSIR2 are properly positioned into a pocket between two β-barrel domains, steric clashes were observed between PsHerA and most α1 helices (4/5) from the dodecameric PsSIR2. This explains the selective binding of the tetradecameric over the dodecameric PsSIR2 by PsHerA.

**Figure 5.**
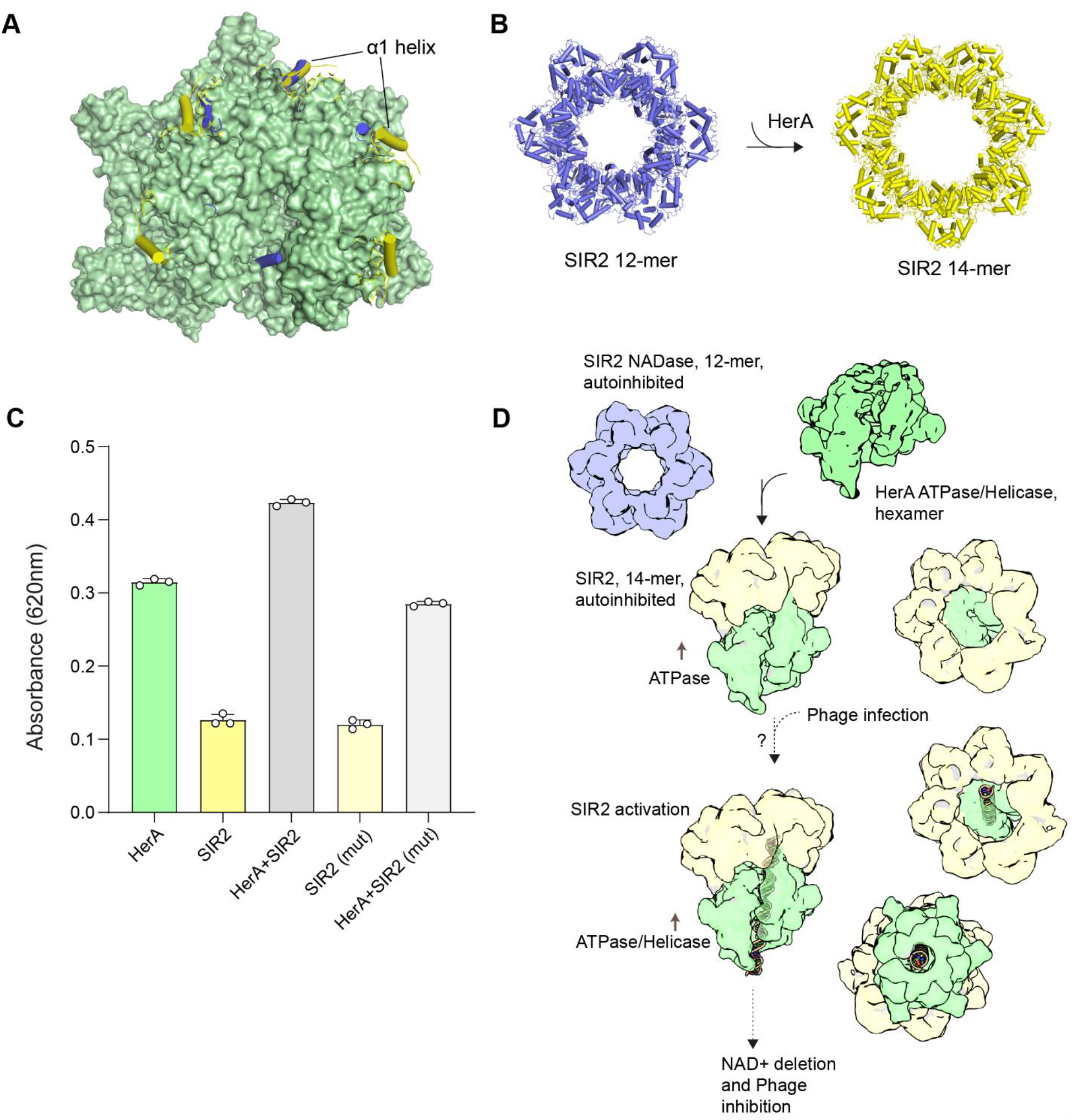
Functional interplay between SIR2 and HerA. (**A**) Superimposition of the dodecameric PsSIR2 and the tetradecameric PsSIR2 in the PsSIR2-HerA complex. The PsHerA hexamer is represented as surface and colored in green. The α1 helices from dodecameric PsSIR2 or tetradecameric PsSIR2 are shown in cartoon representation and colored in slate and yellow, respectively. (**B**) Two assembly states of PsSIR2 were resolved in the PsSIR2-HerA mixture, the dodecameric PsSIR2 is colored in slate and the tetradecameric PsSIR2 is colored in yellow. (**C**) ATPase assays of PsHerA alone or in the presence of PsSIR2. Binding of SIR2 moderately stimulates the ATPase activity of HerA. Three independent measurements were performed. Data are shown as mean ± SEM. (**D**) Proposed schematic for the SIR2-HerA complex in the antiphage defense process.

Intriguingly, the PsSIR2 subunits in the dodecameric and tetradecameric assemblies are nearly identical (Fig. S12A-C), implying that the NADase activity of PsSIR2 is still auto-inhibited in the presence of HerA. Not surprisingly, only a slight increase in NAD^+^ cleavage was observed for the PsSIR2-HerA complex in comparison to PsSIR2 alone (Figure S12D). Consistently, a similar NAD+ degradation profile was observed for PhpSIR2 alone and the PhpSIR2-HerA complex (Figure S12E). Although the assembly interfaces are similar between the dodecameric and tetradecameric PsSIR2, there are outward rotations between subunits to accommodate more subunits in the tetradecameric PsSIR2 (Figure S12C), which particularly weaken the small-small inter-domain interactions, potentially preparing the supramolecular complex for activation by phage-associated molecular patterns upon phage infection. More interestingly, cryo-EM analysis of the PsSIR2-HerA complex sample, however, revealed a significant fraction of tetradecameric PsSIR2 alone in addition to the dodecameric PsSIR2 (Figure 5B and S11), indicating that PsHerA could potentially alter the oligomerization state of PsSIR2 and the tetradecameric PsSIR2 alone may be dissociated from PsHerA.

### The ATPase activity of HerA is stimulated in the presence of SIR2

Superimposition of PsHerA alone and the PsSIR2-HerA complex revealed subtle outward movements of the E and F subunits (Figure S13) around the putative site of ATP hydrolysis in the sequential hydrolysis model (Gates & Martin, 2020). Therefore, PsSIR2 binding is expected to affect PsHerA ATPase activity. Indeed, the ATPase activity of PsHerA was enhanced in the presence of PsSIR2 protein (Figure 5C). By contrast, the PsHerA-binding deficient PsSIR2 mutant did not alter the enzymatic activity of PsHerA. The degradation of ATP has been demonstrated as a strategy of bacterial defense against phages (Rousset *et al*, 2023). Although SIR2-dependent NAD^+^ depletion is proposed to be primarily responsible for phage resistance in SIR2-HerA system (Garb *et al*., 2022), HerA-dependent ATP depletion might also, at least to some extent, contribute to the antiphage activity, particularly in light of the boosted ATPase activity in the context of SIR2-HerA complex.

Together, our studies first revealed the structural basis for the concerted communication between the two proteins in the SIR2-HerA antiphage defense system (Figure 5D). HerA captures the N-terminal α1 helix of SIR2 and prompts the compositional and conformational change of SIR2. At the same time, binding of SIR2 stimulates the ATPase activity of HerA. The phage-associated cue, such as nucleic acids, would be required to be recognized by HerA and sequentially activate the NADase activity of SIR2. Therefore, identification of the phage-associated molecular patterns in the natural context is key to understanding the SIR2-HerA antiphage system in the future.

## Discussion

SIR2 domain-containing proteins are widely distributed in archaea, eukaryotes, and prokaryotes. In comparison to typical archaeal and eukaryotic SIR2 proteins such as *Archaeoglobus fulgidus* sirtuins SIR2-Af1 (Min *et al*., 2001) and yeast Hst2p (Zhao *et al*., 2003) (Figure S4), two insertions (the α8-β3 insertion and the β9-α15 insertion) are present in the large domain of SIR2 proteins from the SIR2-HerA system (Figure 1B and S4A-B). Remarkably, the α8-β3 insertion is involved in the self-assembly of PsSIR2 and PhpSIR2 (Figure 2C and S4D). Similar insertions are also identified in another SIR2-containing protein (ThsA) in the Thoeris bacterial immunity system (Ka *et al*, 2020; Manik *et al*, 2022), which mediates the tetramerization of ThsA (Figure S4C, E). These insertions are therefore likely related to the higher-order assembly of bacterial SIR2 proteins in comparison to archaeal and eukaryotic SIR2 proteins. On the other hand, the zinc finger (Figure S4A-C) and the FEG loop found in archaeal and eukaryotic SIR2 proteins, which are previously suggested to be essential for the deacetylase activity (Avalos *et al*, 2002; Min *et al*., 2001), are absent in bacterial SIR2 proteins, possibly explaining their distinct functions.

The potential binding pocket of NAD^+^ is sterically blocked by the small domain of SIR2 in the apo state. A similar autoinhibition mechanism has also been observed for ThsA in the Thoeris defense system, where the equivalent domain shifts away from the active site upon activation (Manik *et al*., 2022), indicating a conserved regulation mechanism of bacterial SIR2 proteins. (Figure S4C, E). Therefore, conformational movement of the small domain would be necessary to unleash the NADase activity. While no obvious rotation of the small domain with respect to the large domain is seen in the context of the SIR2-HerA complex, the small domain from the adjacent subunit undergoes a movement (Figure S12A-C), potentially compromising the small-small interactions. Thus, our structure may represent a pre-activated state.

Given that the SIR2-dependent antiphage immunity is abrogated when disrupting the large-large inter-domain association (Figure 4D), it is plausible that the large domain-mediated oligomerization of SIR2 might facilitate the activation of the NADase activity. A possible scenario is that the phage-associated signal is transmitted from HerA to the large domain of SIR2, based on the observation that the large domain of SIR2 is responsible for SIR2-HerA complex formation (Figure 4A-B). Coupling diverse enzymes to provide protection from phage appears to be a prevalent strategy for bacterial immunity (Duncan-Lowey *et al*, 2023; Gao *et al*., 2020; Gao *et al*, 2023). HerA ATPase/helicase works in concert with the NurA nuclease in archaea and some bacteria for DNA end resection during the repair of double-stranded DNA breaks (White & Allers, 2018; Yang *et al*, 2023). The toroid-shaped NurA dimer binds to the N-terminal layer of HerA hexamer, unwinding and degrading the DNA translocated by HerA. The working model of the SIR2-HerA system might be analogous to that of the HerA-NurA system. However, for the SIR2-HerA system, the β-barrel domains block the N-terminus of the HerA channel in the split spiral conformation (Figure 3A, 4A). Therefore, translocation of the putative substrate through the HerA channel would likely induce profound rearrangements at the β-barrel domains, thereby possibly leading to the conformational changes of the β-barrel-large domain interface to unleash SIR2 NADase activity. However, further studies would be required to delineate the activation mechanism of the SIR2-HerA complex in the natural context.

## References

Afonine PV, Poon BK, Read RJ, Sobolev OV, Terwilliger TC, Urzhumtsev A, Adams PD (2018) Real-space refinement in PHENIX for cryo-EM and crystallography. Acta Crystallogr D Struct Biol 74: 531–544

Avalos JL, Celic I, Muhammad S, Cosgrove MS, Boeke JD, Wolberger C (2002) Structure of a Sir2 enzyme bound to an acetylated p53 peptide. Mol Cell 10: 523–535

Chen VB, Arendall WB, 3rd, Headd JJ, Keedy DA, Immormino RM, Kapral GJ, Murray LW, Richardson JS, Richardson DC (2010) MolProbity: all-atom structure validation for macromolecular crystallography. Acta Crystallogr D Biol Crystallogr 66: 12–21

Doron S, Melamed S, Ofir G, Leavitt A, Lopatina A, Keren M, Amitai G, Sorek R (2018) Systematic discovery of antiphage defense systems in the microbial pangenome. Science 359

Duncan-Lowey B, Tal N, Johnson AG, Rawson S, Mayer ML, Doron S, Millman A, Melamed S, Fedorenko T, Kacen A et al (2023) Cryo-EM structure of the RADAR supramolecular anti-phage defense complex. Cell 186: 987–998.e915

Dy RL, Richter C, Salmond GP, Fineran PC (2014) Remarkable Mechanisms in Microbes to Resist Phage Infections. Annu Rev Virol 1: 307–331

Emsley P, Lohkamp B, Scott WG, Cowtan K (2010) Features and development of Coot. Acta Crystallogr D Biol Crystallogr 66: 486–501

Fernandez AJ, Berger JM (2021) Mechanisms of hexameric helicases. Critical reviews in biochemistry and molecular biology 56: 621–639

Gao L, Altae-Tran H, Böhning F, Makarova KS, Segel M, Schmid-Burgk JL, Koob J, Wolf YI, Koonin EV, Zhang F (2020) Diverse enzymatic activities mediate antiviral immunity in prokaryotes. Science 369: 1077–1084

Gao Y, Luo X, Li P, Li Z, Ye F, Liu S, Gao P (2023) Molecular basis of RADAR anti-phage supramolecular assemblies. Cell 186: 999–1012.e1020

Garb J, Lopatina A, Bernheim A, Zaremba M, Siksnys V, Melamed S, Leavitt A, Millman A, Amitai G, Sorek R (2022) Multiple phage resistance systems inhibit infection via SIR2-dependent NAD+ depletion. Nature microbiology 7: 1849–1856

Gates SN, Martin A (2020) Stairway to translocation: AAA+ motor structures reveal the mechanisms of ATP-dependent substrate translocation. Protein Science 29: 407–419

Jumper J, Evans R, Pritzel A, Green T, Figurnov M, Ronneberger O, Tunyasuvunakool K, Bates R, Žídek A, Potapenko A et al (2021) Highly accurate protein structure prediction with AlphaFold. Nature 596: 583–589

Ka D, Oh H, Park E, Kim JH, Bae E (2020) Structural and functional evidence of bacterial antiphage protection by Thoeris defense system via NAD(+) degradation. Nat Commun 11: 2816

Koopal B, Potocnik A, Mutte SK, Aparicio-Maldonado C, Lindhoud S, Vervoort JJ, Brouns SJ, Swarts DC (2022) Short prokaryotic Argonaute systems trigger cell death upon detection of invading DNA. Cell 185: 1471–1486. e1419

Labrie SJ, Samson JE, Moineau S (2010) Bacteriophage resistance mechanisms. Nat Rev Microbiol 8: 317–327

Li Z, Kaur P, Lo C-Y, Chopra N, Smith J, Wang H, Gao Y (2022) Structural and dynamic basis of DNA capture and translocation by mitochondrial Twinkle helicase. Nucleic acids research 50: 11965–11978

Liebschner D, Afonine PV, Baker ML, Bunkóczi G, Chen VB, Croll TI, Hintze B, Hung LW, Jain S, McCoy AJ et al (2019) Macromolecular structure determination using X-rays, neutrons and electrons: recent developments in Phenix. Acta Crystallogr D Struct Biol 75: 861–877

Makarova KS, Wolf YI, Koonin EV (2013) Comparative genomics of defense systems in archaea and bacteria. Nucleic Acids Res 41: 4360–4377

Makarova KS, Wolf YI, Snir S, Koonin EV (2011) Defense islands in bacterial and archaeal genomes and prediction of novel defense systems. J Bacteriol 193: 6039–6056

Makarova KS, Wolf YI, van der Oost J, Koonin EV (2009) Prokaryotic homologs of Argonaute proteins are predicted to function as key components of a novel system of defense against mobile genetic elements. Biol Direct 4: 29

Manik MK, Shi Y, Li S, Zaydman MA, Damaraju N, Eastman S, Smith TG, Gu W, Masic V, Mosaiab T et al (2022) Cyclic ADP ribose isomers: Production, chemical structures, and immune signaling. Science 377: eadc8969

Millman A, Melamed S, Leavitt A, Doron S, Bernheim A, Hör J, Garb J, Bechon N, Brandis A, Lopatina A et al (2022) An expanded arsenal of immune systems that protect bacteria from phages. Cell Host Microbe 30: 1556–1569.e1555

Min J, Landry J, Sternglanz R, Xu R-M (2001) Crystal structure of a SIR2 homolog–NAD complex. Cell 105: 269–279

Mirdita M, Schütze K, Moriwaki Y, Heo L, Ovchinnikov S, Steinegger M (2022) ColabFold: making protein folding accessible to all. Nature methods 19: 679–682

Neuwald AF, Aravind L, Spouge JL, Koonin EV (1999) AAA+: A class of chaperone-like ATPases associated with the assembly, operation, and disassembly of protein complexes. Genome research 9: 27–43

North BJ, Verdin E (2004) Sirtuins: Sir2-related NAD-dependent protein deacetylases. Genome biology 5: 1–12

Ogura T, Wilkinson AJ (2001) AAA+ superfamily ATPases: common structure–diverse function. Genes to Cells 6: 575–597

Pettersen EF, Goddard TD, Huang CC, Meng EC, Couch GS, Croll TI, Morris JH, Ferrin TE (2021) UCSF ChimeraX: Structure visualization for researchers, educators, and developers. Protein Sci 30: 70–82

Puchades C, Sandate CR, Lander GC (2020) The molecular principles governing the activity and functional diversity of AAA+ proteins. Nature Reviews Molecular Cell Biology 21: 43–58

Punjani A, Rubinstein JL, Fleet DJ, Brubaker MA (2017) cryoSPARC: algorithms for rapid unsupervised cryo-EM structure determination. Nat Methods 14: 290–296

Rousset F, Yirmiya E, Nesher S, Brandis A, Mehlman T, Itkin M, Malitsky S, Millman A, Melamed S, Sorek R (2023) A conserved family of immune effectors cleaves cellular ATP upon viral infection. Cell 186: 3619–3631.e3613

Rzechorzek NJ, Blackwood JK, Bray SM, Maman JD, Pellegrini L, Robinson NP (2014) Structure of the hexameric HerA ATPase reveals a mechanism of translocation-coupled DNA-end processing in archaea. Nature communications 5: 5506

Schuck P (2000) Size-distribution analysis of macromolecules by sedimentation velocity ultracentrifugation and lamm equation modeling. Biophysical journal 78: 1606–1619

Stern A, Sorek R (2011) The phage-host arms race: shaping the evolution of microbes. Bioessays 33: 43–51

Thompson RF, Iadanza MG, Hesketh EL, Rawson S, Ranson NA (2019) Collection, preprocessing and on-the-fly analysis of data for high-resolution, single-particle cryo-electron microscopy. Nat Protoc 14: 100–118

White MF, Allers T (2018) DNA repair in the archaea-an emerging picture. FEMS Microbiol Rev 42: 514–526

White SR, Lauring B (2007) AAA+ ATPases: achieving diversity of function with conserved machinery. Traffic 8: 1657–1667

Yang J, Sun Y, Wang Y, Hao W, Cheng K (2023) Structural and DNA end resection study of the bacterial NurA-HerA complex. BMC Biol 21: 42

Zaremba M, Dakineviciene D, Golovinas E, Zagorskaitė E, Stankunas E, Lopatina A, Sorek R, Manakova E, Ruksenaite A, Silanskas A et al (2022) Short prokaryotic Argonautes provide defence against incoming mobile genetic elements through NAD(+) depletion. Nat Microbiol 7: 1857–1869

Zhang K (2016) Gctf: Real-time CTF determination and correction. J Struct Biol 193: 1–12

Zhao K, Chai X, Clements A, Marmorstein R (2003) Structure and autoregulation of the yeast Hst2 homolog of Sir2. Nature Structural & Molecular Biology 10: 864–871

Zheng SQ, Palovcak E, Armache JP, Verba KA, Cheng Y, Agard DA (2017) MotionCor2: anisotropic correction of beam-induced motion for improved cryo-electron microscopy. Nat Methods 14: 331–332

Zivanov J, Nakane T, Forsberg BO, Kimanius D, Hagen WJ, Lindahl E, Scheres SH (2018) New tools for automated high-resolution cryo-EM structure determination in RELION-3. Elife 7

